# Dynamic competition between bottom-up saliency and top-down goals in early visual cortex

**DOI:** 10.1101/2025.08.22.671530

**Authors:** Dan Wang, Kabir Arora, Jan Theeuwes, Stefan Van der Stigchel, Surya Gayet, Samson Chota

**Author notes:** **Corresponding author:** Dan Wang, Heidelberglaan 1, 3584 CS, Utrecht, Netherlands, **Email:**. Surya Gayet and Samson Chota contributed equally to this work. Jan Theeuwes was supported by a European Research Council (ERC) advanced grant 833029 – [LEARNATTEND] and by a NWO Open competition grant 25 406.21.GO.034.

## Abstract

Task-irrelevant yet salient stimuli can elicit automatic, bottom-up attentional capture and compete with top-down, goal-directed processes for neural representation. However, the temporal dynamics underlying this competition, and how they influence early visual processing, remain poorly understood. Here, we combined electroencephalography (EEG) with Rapid Invisible Frequency Tagging (RIFT) to non-invasively and simultaneously track early visual cortex responses to target and distractor. Both target and distractor evoked stronger initial RIFT responses than nontargets, reflecting top-down and bottom-up attentional effects on early visual processing. Importantly, the presence of a distractor attenuated the initial RIFT response to the target, reflecting competition during the initial stages of visual processing and predicting subsequent behavioral performance. RIFT responses to the distractor eventually even decreased below responses to the target and nontarget, representing active suppression of task-irrelevant but salient stimuli. We show that the dynamic interplay between top-down control and bottom-up saliency directly impacts early visual responses, thereby illuminating a complete timeline of attentional competition in visual cortex.

## Introduction

Imagine driving down a busy road, focusing on the surrounding traffic, when a flashing billboard suddenly catches your eye and briefly distracts you from the roadway ahead. This illustrates how attentional control arises from the interaction between two competing processes: bottom-up control driven by saliency (e.g., the flashing billboard) whereby attention is automatically captured by elements that stand out from the environment (Theeuwes, 1991; 1992; 2021), and top-down control (e.g., maintaining focus on the roadway) which directs attention based on goals and intentions (Folk and Remington, 1998). It is widely accepted that both processes contribute to attentional selection (Luck et al., 2021). According to the biased competition framework (Desimone & Duncan, 1995; Luck et al., 1997; Tsotsos et al., 1995) objects in the visual field compete for neural representation in visual cortex. This competition is initially driven by bottom-up salience during the early feedforward sweep of sensory processing and is subsequently shaped by top-down signals, likely conveyed via feedback connections from higher-level cortical areas (Beck & Kastner, 2009; Theeuwes, 2010). It remains unclear, however, how these processes unfold over time within early visual cortex. Here, we test whether initial bottom-up salience signals and subsequent top-down control mechanisms are both reflected in early visual cortex responses to competing stimuli.

To determine how top-down and bottom-up processes unfold over time in early visual cortex, we employed Rapid Invisible Frequency Tagging (RIFT) while participants performed the additional singleton task (Theeuwes, 1991; 1992; 2010). In this task, participants search for a shape singleton target among nontarget items (e.g., a green diamond among green circles). On some trials, one of the nontarget items is a salient but irrelevant color singleton distractor (e.g., a red circle). Typically, response times increase on “distractor present” trials compared to “distractor absent” trials, indicating that the distractor captured attention in a bottom-up way. By utilizing RIFT, we are able to track, in time, how biased competition unfolds between the bottom-up salience of the distractor and the top-down relevance of the target in early visual cortex. Specifically, we can test (1) whether the presence of a salient task-irrelevant distractor reduces early visual cortex responses to a concurrent task-relevant target, and (2) whether subsequent top-down control mechanisms further reduce responses to a salient but task-irrelevant distractor.

RIFT works by modulating the luminance of one or more visual stimuli at distinct high frequencies (e.g., 60 Hz and 64 Hz), which elicits frequency-matching periodic activity in the EEG signal originating from early sensory areas (V1/V2; Arora et al., 2025; Dietz et al., 2025 (preprint); Duecker et al., 2021; 2025; Ferrante et al., 2023; Minarik et al., 2023; Seijdel et al., 2023; Zhigalov et al., 2019). These periodic responses enable highly time-resolved and spatially specific tracking of attention in the early visual cortex (Arora et al., 2025; Duecker et al., 2025; Ferrante et al., 2023; Zhigalov et al., 2019). Importantly, because stimulus luminance is modulated at frequencies far above the critical flicker fusion threshold (Landis, 1954; e.g., ∼40 Hz), the flicker is imperceptible to observers and does not perceptually interfere with the ongoing task (Spaak et al., 2024). Together, these properties make RIFT a powerful tool for disentangling visually evoked responses to concurrently presented visual events. Here, this technique allows us to test whether—and how—the competition between target and distractor stimuli unfolds in early visual cortex.

## Results

### Behavioral results

To evaluate whether the presence of distractor affected behavioral performance, we conducted a paired-sample t-test comparing mean response times (RTs) for correct trials between the distractor present and absent conditions. This revealed that RTs were significantly slower when the distractor was present (*Mean* = 924 ms) compared to when it was absent (*Mean* = 873 ms, *t*(23) = 8.582, *p* < 0.001, Cohens’ *d* = 1.752; Figure 1B), a finding typically interpreted as evidence that the salient distractor captured attention. A similar effect was observed for accuracy (see supplementary materials, Figure S1).

**Figure 1.**
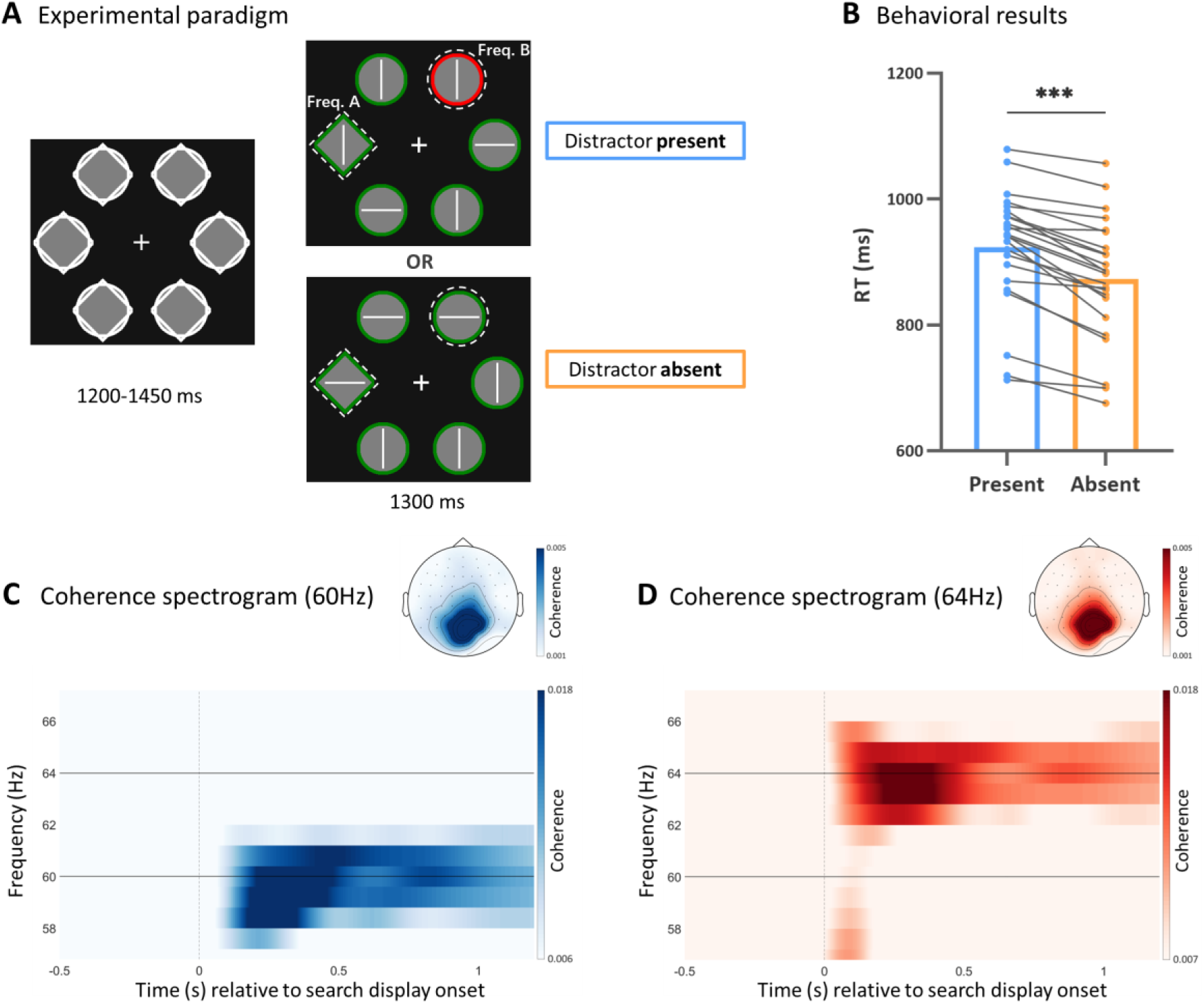
**(A)** Experimental paradigm. After an initial display consisting of placeholders, the search display was presented. Participants were instructed to search for a unique shape singleton target (here a diamond among circles) and respond as quickly and accurately as possible to orientation of the line segment inside it. On half of the trials, a color distractor was present (here: red among green). Target and distractor (or one of the nontargets) were frequency tagged (luminance-modulated) at 60 Hz and 64 Hz throughout the search display (see Tagging manipulation for details). Note: the figure is not to scale; no outlines were visible around the flickering regions in the actual experiment. **(B)** Behavioral results. Participants were slower to find the target when a distractor was present. The blue bar represents reaction times in the distractor present condition, while the orange bar represents the distractor absent condition. Each dot indicates the mean response time of an individual participant. \*\*\**p* < 0.001. **(C)** Time-frequency plot of coherence (phase realigned 60 Hz), averaged across participants and individuals top 6 channels. *Top right inset:* Scalp topography of average 60 Hz coherence across the 1.2-second after search display onset. **(D)** Time-frequency plot of coherence (64 Hz), averaged across participants and individuals top 6 channels. *Top right inset:* Scalp topography of average 64 Hz coherence across the 1.2-second after search display onset.

### Validation of frequency-specific neural responses

We verified whether our frequency-tagging manipulation successfully elicited corresponding frequency-specific neural responses by calculating the coherence between the EEG signal and the corresponding tagging frequencies. Because we randomly shifted the phase of the 60 Hz tag relative to the 64 Hz tag, we computed coherence spectrograms separately for each frequency (see Methods section). The resulting spectrograms showed clear peaks at 60 Hz (Figure 1C) and 64 Hz (Figure 1D) following flicker onset, with strongest responses over parietal and occipital electrodes (see topography insets, individual traces in supplementary, Figure S2; S3), confirming successful retrieval of the tagging signals from the EEG.

### RIFT responses to the distractor and nontarget

We tested whether attentional capture by the distractor was reflected in the RIFT responses. Compared to the nontarget, the distractor evoked significantly stronger coherence in an initial time window (*p* = 0.036; [cluster extent: ∼150 to ∼350 ms]; cluster-based permutation test) and significantly weaker coherence in a later time window (*p* = 0.019; [cluster extent: ∼640 to ∼900 ms]; cluster-based permutation test; Figure 2A, left). To further characterize the temporal dynamics of the visual processing at the early time window (0-600 ms), we conducted a one-sided t-test to examine whether the time-to-peak of the coherence traces differed between distractor and nontarget. However, the time-to-peak of the coherence trace for the distractor was not significantly different from that for the nontarget (*p* = 0.103, *t*(23) = 1.302, Cohen’s *d* = 0.266; Figure 2A, right).

**Figure 2.**
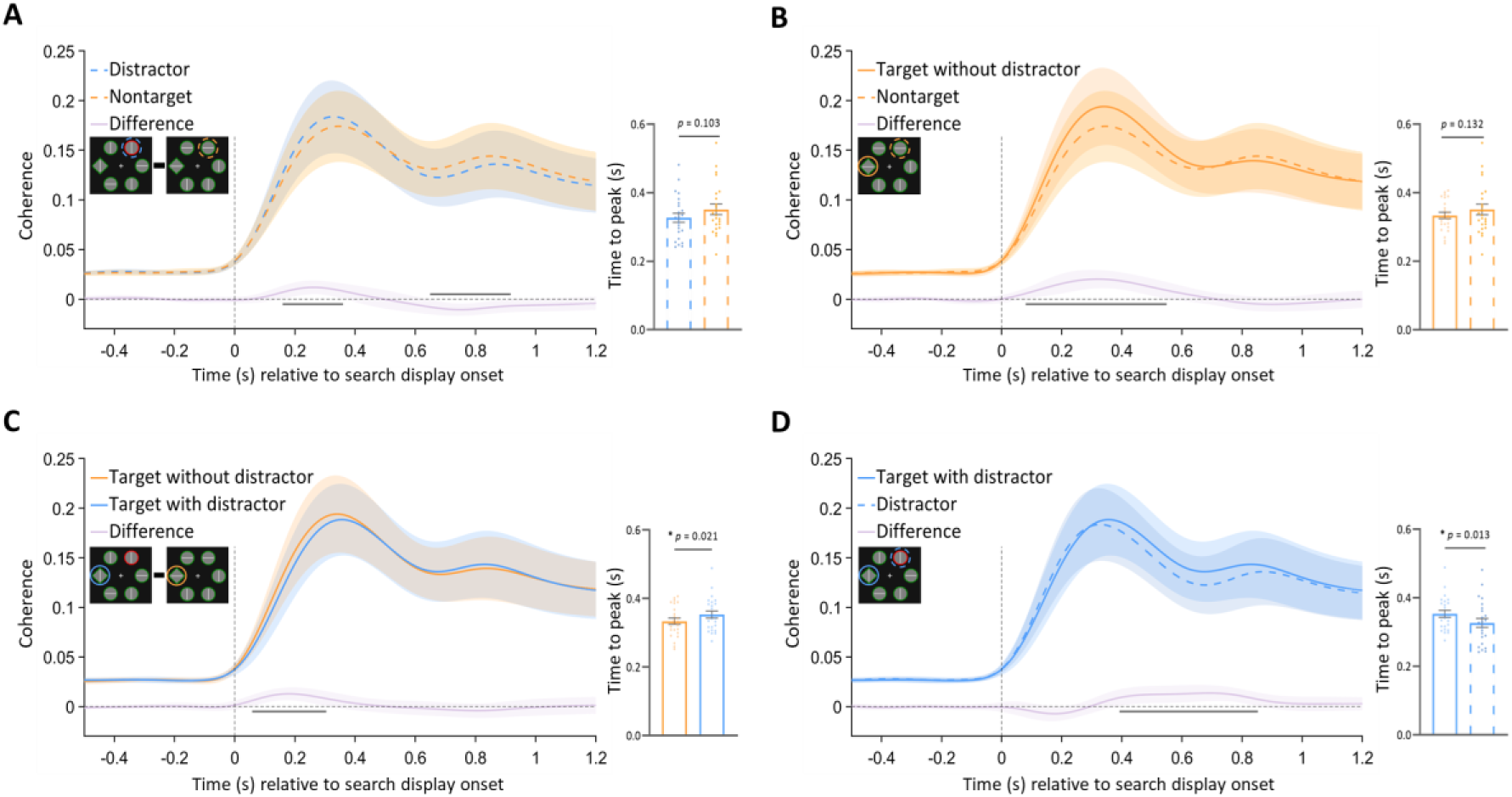
RIFT responses across experimental conditions. **(A).** Coherence time-series of distractor (blue dashed), nontarget (orange dashed) and the difference (purple solid). Shaded areas represent 95% confidence intervals of the mean. Significant clusters (from cluster based-permutation tests) are indicated by horizontal solid black lines. Right bar graphs show the time-to-peak analysis of the coherence trace (0-600 ms) for each condition. **(B).** Coherence time-series of target without distractor (orange solid), nontarget (orange dashed) and the difference (purple solid). **(C).** Coherence time-series of target with distractor (blue solid), target without distractor (orange solid) and the difference (purple solid). **(D).** Coherence time-series of target with distractor (blue solid), distractor (blue dashed) and the difference (purple solid).

Coherence measures thus indicated that more attention was allocated to the salient distractor than to the nontarget items initially, as indexed by a stronger stimulus-specific RIFT responses, signifying early attentional capture. Critically, attention to the salient distractor later fell below that directed to the nontarget items, reflecting attentional disengagement and suppression. This pattern compellingly illustrates the temporal unfolding of attentional allocation in early visual cortex, revealing how attention is first captured by the salient distractor and subsequently withdrawn and even suppressed over time.

### RIFT responses to target and nontarget

To examine whether attentional selection of the target was reflected in the RIFT responses when no distractor was present, we compared target-evoked coherence with the coherence evoked by the nontarget within the same trial. Target-related coherence was significantly stronger than coherence for nontarget in an initial time window (*p* = 0.0001; [cluster extent: ∼75 to ∼540 ms]; cluster-based permutation test; Figure 2B, left). The time-to-peak of the coherence trace did not differ significantly between the two during the early time period (0-600 ms; *p* = 0.132, *t*(23) = 1.145, Cohen’s *d* = 0.234; Figure 2B, right). These results suggest that when no salient competitor is present, attention is initially allocated to the only salient item in the display, allowing for fast and accurate selection of the salient target.

### RIFT responses to target with and without distractor

We tested whether the presence of a distractor affected early visual processing of the target. To this end, we compared coherence evoked by the target in the presence versus absence of a distractor. Coherence was significantly higher when the distractor was absent in an initial time window (*p* = 0.015; [cluster extent: ∼50 to ∼300 ms]; cluster-based permutation test; Figure 2C, left). Furthermore, the coherence for the target reached its peak earlier in the early time window (0-600 ms) when the distractor was absent compared to when it was present (*p* = 0.021, *t*(23) = 2.149, Cohen’s *d* = 0.439; Figure 2C, right). These results demonstrate that the presence of a distractor results in less attention being allocated to the target, consistent with the notion of biased competition (Desimone & Duncan, 1995).

### RIFT responses to targets and distractor

To investigate the dynamics of attentional competition between the target and the distractor, we statistically compared their coherence traces in the distractor present condition. No significant differences in coherence were observed between the target and the distractor at the beginning of the trial (Figure 2D, left). However, within the early time window (0–600 ms), the peak coherence occurred significantly later for the target than for the distractor (*p* = 0.013, *t*(23) = 2.395, Cohen’s *d* = 0.489; Figure 2D, right). Critically, target-evoked coherence significantly exceeded distractor-evoked coherence in a later time window (*p* = 0.002; [cluster extent: ∼390 to ∼840 ms]; cluster-based permutation; Figure 2D, left).

Taken together, these findings suggest that the target and distractor are in direct competition for attentional resources during the early stage of processing, with the distractor initially capturing attention and thereby delaying visual processing of the target. Over time, the RIFT responses to the distractor were attenuated and even suppressed relative to nontarget items, thereby resolving the competition between the distractor and target, enabling the selection of the target. These results highlight that RIFT is well-suited to measure changes in neuronal excitability in early visual cortex associated with attentional competition during visual search.

### Correlation between RIFT responses and behavioral RTs

To investigate whether the measured RIFT responses to target and distractor were related to participants’ behavior, we computed trial-wise correlations between Trial-Ensemble Phase Similarity (TEPS, a phase-based, single-trial measure of the tagging signal; see Methods section) and RTs. Because we were specifically interested in how the competition between the target and the distractor was resolved over time, we calculated the difference in RIFT responses between the target and distractor within the same trial and correlated this differential RIFT response with the RTs to the target across trials (Figure 3). We observed a significant negative correlation between TEPS and RTs in an initial time window (*p* = 0.003; [cluster extent: ∼290 to ∼520 ms]; cluster-based permutation; Figure 3), indicating that the greater the RIFT responses to the target compared to the distractor, the faster the participant responded to the target. This finding demonstrates that the RIFT responses capture behaviorally relevant processes.

**Figure 3.**
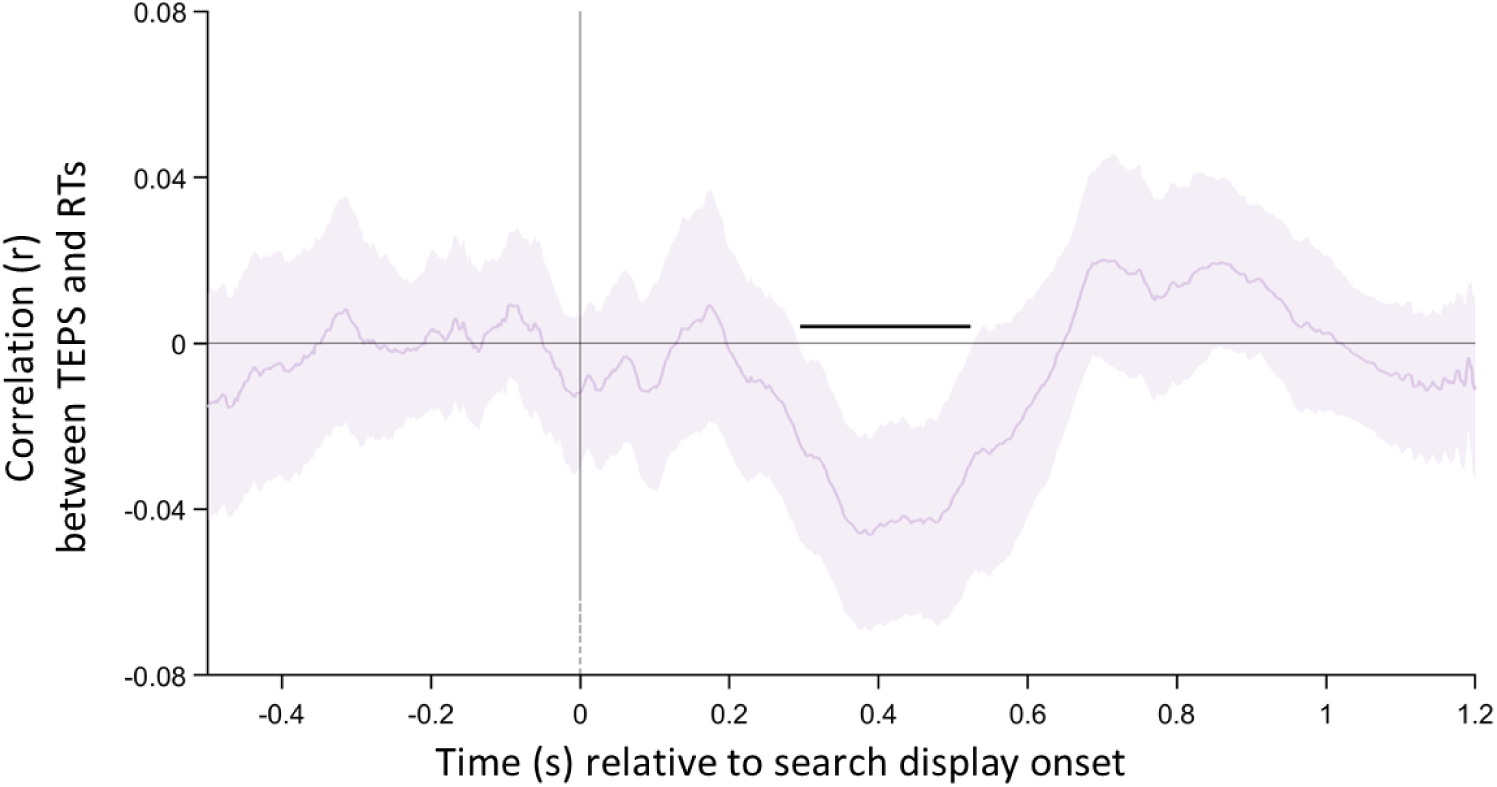
Time-resolved, trial-wise correlation between RTs and the TEPS difference between target and distractor within the same trial. Only correct trials in the distractor present condition used. Shaded areas represent 95% confidence intervals of the mean. Significant clusters (from cluster based-permutation tests) are indicated by horizontal solid black lines.

## Discussion

The present study employed Rapid Invisible Frequency Tagging (RIFT) with EEG to examine how competition between top-down attention (to the target) and bottom-up attention (to the distractor) unfolds over time in visual cortex. In line with the biased competition framework (Desimone & Duncan, 1995; Reynolds & Desimone, 2003), the results indicate that early in processing the distractor briefly dominates the competition, whereas at later stages the target prevails. Crucially, the outcome of this competition is a clear neural representation in visual cortex of the selected (winning) object, accompanied by a diminished representation of the non-selected (losing) objects.

In the current task, participants correctly responded to the target well above chance level, indicating that ultimately the target is selected and wins the competition. However, the strength of this study lies in how the RIFT responses to both the target and distractor jointly provide insight into how this competition is resolved, up to the point of the eventual selection of the target. It is evident that in distractor-absent trials, there is basically no competition (the target is the only salient element in the display) and the RIFT responses show that in this condition target processing dominates from the earliest moment onwards, giving rise to fast and accurate responses.

However, due to the limited processing capacity of the visual system, competition arises when both the target and distractor are simultaneously present, both competing for neural representation. The results show that, early on—during the initial feedforward sweep of sensory processing—the competition is won by the salient distractor: RIFT responses to the target were delayed, while responses to the distractor exceeded those elicited by nontarget items. Later in time, RIFT responses to the salient distractor dropped not only below responses to the target, but also below responses to nontarget items. This suggests that salient but task-irrelevant objects in the environment not only cease to attract attention over time but may also become suppressed to facilitate the neural representation of the task-relevant objects. Critically, within-subject, trial-wise correlations reveal that larger RIFT responses to target compared with distractor (a direct measure of competition in the early visual cortex) are associated with faster reaction times to find the target, highlighting the functional role of these early sensory modulations for the successful completion of goal-directed behavior.

Overall, the pattern of RIFT responses is consistent with stimulus-driven accounts of perceptual competition, which propose that during the initial feedforward sweep of visual processing attention is automatically captured by the most salient element in the display (Theeuwes, 1992; 2010; 2025). Only later, through feedback signals, top-down processing allows attention to be disengaged from the distractor (Theeuwes et al., 2000). The below-baseline RIFT response of the distractor indicates that disengagement even involves suppression, in line with the notion of reactive suppression (Geng, 2014).

A recent study by Klink et al (2023) provides similar evidence supporting initial capture followed by rapid disengagement. In their research, Klink and colleagues examined neural responses in area V4 of macaque monkeys performing an eye movement-based version of the same paradigm that was used here (i.e., the additional singleton paradigm). Eye-tracking data suggested that the salient distractor was effectively ignored, as the monkeys’ eyes moved directly to the target. Yet, the neuronal activity of V4 neurons showed a different picture: Early on, during the initial stage, there was attentional enhancement at the location occupied by the salient distractor. This initial enhancement was followed by suppression, occurring about 150 ms later. These data show that even though behaviorally there appears to be successful inhibition of the salient distractor, this inhibition was preceded by attentional capture, providing evidence for the fast disengagement hypothesis (Theeuwes et al., 2000). Thus, the pattern of neuronal activity in V4 neurons reported by Klink et al. (2023) is consistent with the current pattern of findings in early visual cortex, obtained non-invasively in human subjects.

Also using the additional singleton paradigm, Lin et al (2024) recently recorded human intracranial signals covering multiple brain regions (but with very limited coverage in early visual cortex). They were able to dissociate distractor-specific representations from target signals in the high-frequency range (60–100 Hz). Consistent with the current findings, Lin et al. found that initially salient distractors were processed around 220 ms after stimulus onset, while at the same time, target-related processing was attenuated. Their findings highlight the competition for neural representation between target and distractor, consistent with the biased competition framework (Desimone & Duncan, 1995; Reynolds & Desimone, 2003). The present work extends the findings of Lin and colleagues (non-invasively), by specifically revealing that biased competition between target and distractor stimuli even influences early visual cortex responses.

Using the same paradigm, previous studies have recorded scalp EEG to examine shifts of attention toward targets and distractors (e.g., Hickey et al., 2006; 2010; Schubö, 2009; Wang et al., 2019). Specifically, the N2pc component of the event-related potential was used to track the allocation of attention to lateralized positions in the search array. For example, in Experiment 2 of Hickey et al. (2006), both the distractor and the target (on different trials) elicited an N2pc when they appeared on opposite sides of the array. Critically, however, the pattern of N2pc responses indicated that attention was initially captured by the salient distractor before shifting to the target. This is consistent with a stimulus-driven account of attentional capture (Theeuwes, 1992; 2010) and with our current results. Importantly, however, our approach extends prior work by simultaneously measuring neural responses to both the target and the salient distractor within the same trial. This allowed us to directly assess their competitive interaction in visual cortex and link these neural dynamics to behavioral performance.

The temporal profile of RIFT responses suggests that suppression of the distractor relative to the nontarget items might be necessary for successful processing of the target. In other words, consistent with the biased competition framework, the competition between the distractor and the target must be resolved in favor of the target to enable its processing. Similar findings have been reported previously (Lin et al., 2024; Klink et al., 2023; Duecker et al., 2025; Cosman et al., 2018; Forschack et al., 2022). As previously argued (Theeuwes, 2010; Born et al., 2011) distractor suppression may be necessary for attentional disengagement and is likely driven by top-down control signals originating from higher-order cortical regions such as the inferior frontal gyrus, prefrontal cortex, and area V4. These regions have been implicated in facilitating attentional shifts away from distractors and toward goal-relevant stimuli (Cosman et al., 2018; Klink et al., 2023; de Fockert & Theeuwes, 2012).

One potential concern is that differences in eye movements toward the tagged stimuli might influence RIFT responses, since neural responses tend to be stronger for stimuli presented near the fovea (Wandell et al., 2007). To address this concern, we removed trials in which fixation was not adequately maintained. In addition, trial-wise correlation analyses between gaze bias and RIFT responses for both targets and salient distractors revealed no significant correlations (see Supplementary, Figure S5), suggesting that gaze position did not predict RIFT responses. Moreover, previous studies have shown that RIFT responses are not affected by small eye movements around fixation (Arora et al., 2025; Duecker et al., 2025), which further supports the conclusion that our findings reflect attentional process rather than fixation instability or gaze shifts.

We interpret the changes in RIFT-evoked neuronal excitability as reflecting changes in the responsiveness of early visual cortex. Consistent with this view, previous studies have taken RIFT response modulations to indicate changes in neural excitability within early visual areas, including both primary and secondary regions (Duecker et al., 2021; 2025). Supporting this interpretation, studies combining RIFT with magnetoencephalography (MEG) have consistently localized RIFT responses to early visual areas V1 and V2 (Duecker et al., 2021; 2025; Minarik et al., 2023; Schneider et al., 2023; Zhigalov & Jensen, 2020). Taken together, this body of evidence provides a strong basis for linking our observed RIFT responses to early visual cortex activity. The present findings demonstrate that both top-down and bottom-up factors shape even the earliest stages of visual processing when stimuli compete for representation.

In summary, the present study demonstrates that, during visual search, a salient distractor initially dominates processing in early visual cortex, reflecting a strong bottom-up drive. Subsequently, reactive suppression of the distractor is accompanied by a relative enhancement of the neural response to the target, which may jointly underlie successful target selection. Using RIFT with EEG, the current study reveals the dynamic interplay between bottom-up salience and top-down control in resolving attentional competition within early visual cortex.

## Methods

### Participants

To determine the appropriate sample size, we conducted a priori power analysis using G*Power 3.1 (Faul et al., 2007). Assuming a paired-samples t-test, an alpha level of 0.05, an effect size (dz) of 0.6, and a desired power of 0.8, the analysis indicated that 24 participants would be required. This sample size is comparable to those used in prior studies on attentional capture with EEG (e.g., Hickey et al., 2006; 2010; with 18 participants) and rhythmic sensory stimulation (e.g., Arora et al., 2025; Duncan et al., 2025; both with 24 participants). Four participants were replaced: three due to an excessive proportion of saccades (93.4%, 63.9%, and 52.8%, respectively), one due to below-chance search task performance (48% accuracy). The final sample thus consisted of 24 participants (*mean* age = 22.83 years, *SD* = 2.87; 22 females). All participants had normal or corrected-to-normal vision and reported no history of epilepsy or cognitive impairments. Written informed consent was obtained prior to participation, and participants received either monetary compensation or course credit. The study was approved by the Ethics Committee of Utrecht University.

### Apparatus

Stimuli were presented using a ProPixx projector (VPixx Technologies Inc., QC, Canada; 960 × 540 pixels, 480 Hz refresh rate) in a rear-projection format (screen size: 48 × 27.2 cm). All stimuli were created using MATLAB 2021 (The MathWorks, Inc.) with the PsychToolbox extension (Kleiner et al., 2007). The viewing distance was maintained at 72 cm using a chin and forehead rest. Gaze was tracked using an EyeLink SR (SR Research, Ontario, Canada) eye tracker, which recorded data from both eyes at a sampling rate of 500 Hz.

EEG data were recorded using a 64-channel ActiveTwo BioSemi system (BioSemi B.V., Amsterdam, The Netherlands) at a sampling rate of 2048 Hz. To monitor eye movements and detect ocular artifacts, two additional electrodes were placed: one above the left eye to record vertical eye movements and one on the outer canthus of the left eye to capture horizontal eye movements. Before the experiment, signal quality across all channels was assessed and optimized using BioSemi ActiView software, ensuring stable and high-quality recordings.

### Procedure

In the main experiment (depicted in Figure 1A), participants were instructed to maintain their gaze fixation on the central cross throughout the entire experiment. Each trial began with the presentation of a placeholder display for a randomly varying duration between 1200 and 1450 ms. Following the placeholder display, a search display appeared for a fixed duration of 1300 ms. Participants were instructed to identify whether the line segment inside the unique (target) shape (circle or diamond) was vertical (press “P”) or horizontal (press “Q”) as quickly and accurately as possible with left and right index finger respectively. Upon participant response, the color of the central cross changed from white to black. Participants completed 40 practice trials to familiarize the experimental procedure. The main experiment consisted of 1152 trials, divided into 8 blocks, and lasted approximately one hour. Prior to the experiment, a 9-point calibration was conducted to ensure accurate gaze measurements. This calibration was repeated after every two experimental blocks to maintain gaze tracking precision throughout the session.

### Stimuli

All stimuli were displayed on a uniform dark background with an RGB value of (20, 20, 20). The placeholder display comprised six shapes, each formed by superimposing a diamond (4.3° × 4.3° square rotated 45°) onto a circle (radius = 2.1°). The sizes of the circle and diamond matched those of the stimuli used in the subsequent search task. The outline of each shape was white (RGB: 255, 255, 255), while the inner area was filled with a mid-gray color (RGB: 127.5, 127.5, 127.5). These six shapes were evenly spaced along an imaginary circle (radius = 6°) centered around the fixation cross (1.2° in length; RGB: 255, 255, 255).

In the search task, six items were displayed in the same spatial arrangement as in the placeholder display, ensuring spatial consistency throughout the experiment. In the *distractor present* condition, the array consisted of one shape singleton target, one color singleton (salient) distractor, and four nontargets. In the *distractor absent* condition, the array contained one shape singleton target and five nontargets. The nontargets always shared the same color as the target and the same shape as the distractor. The target was either a circle or a diamond. When the target was a circle, all distractors were diamonds and vice versa. The outline color of the target was either green (RGB: 0, 131, 0) and that of the color-salient distractor was red (RGB: 255, 0, 0), or vice versa. By varying both the shape and the color assignment across trials, participants could not proactively prepare for target or distractor features before the onset of the search array. Each item in the search array contained a central white line segment (3.1° in length), oriented either horizontally or vertically. The inner area of the target and one distractor (salient or nontarget, depending on condition) was luminance-modulated (excluding the white line segment) at either 64 Hz or 60 Hz, making them perceptually indistinguishable from mid-gray (Arora et al., 2025). The inner areas of the remaining nontargets were filled with mid-gray. The target appeared randomly at one of the six positions with equal probability, with the constraint that the distractor was never placed directly adjacent to the target.

### Tagging manipulation

We implemented RIFT stimulation from specific spatial locations in the visual field (corresponding to the target and distractor stimuli) by sinusoidally modulating the luminance of the inner area of stimuli at high temporal frequency (Arora et al., 2025; Drijvers et al., 2021). Tagging was applied throughout the entire duration of the search display. The two tagging frequencies (60Hz and 64Hz) were counterbalanced across target and distractor stimuli to ensure that any difference in RIFT responses between stimuli could not be attributed to differences between tagging frequencies. In the distractor present condition, the target was tagged with one frequency (either 60 Hz or 64 Hz) and the distractor with the other. In the distractor absent condition, the target was tagged with one frequency and one of the nontargets with the other.

To improve the temporal resolution of the RIFT responses given the inherent trade-off between time and frequency resolution (i.e., the Heisenberg uncertainty principle for signals), we implemented a phase randomization procedure as described below. The 64 Hz tagging signal was phase-locked to the onset of the search display in all trials. In contrast, the phase of the 60 Hz tagging signal was randomized at one of eight equally spaced phase offsets within a cycle (excluding 0° to avoid overlap with the 64 Hz component), randomly assigned on each trial. These phase offsets were recorded and later used to reconstruct a phase-aligned EEG signal for subsequent analysis (see RIFT responses section). Decoupling the 60 Hz and 64 Hz signals based on phase facilitates the separation of tags during EEG analysis, specifically when using methods that quantify the degree of phase alignment across trials (i.e. coherence, see RIFT responses section). Practically, it enables the use of broader bandpass filters when isolating the tagged responses, thereby enhancing temporal resolution without compromising frequency specificity.

### EEG pre-processing

All data analysis was conducted in MATLAB using the Fieldtrip toolbox (Oostenveld et al., 2011). The EEG data was first re-referenced to the average of all channels (excluding poor channels determined by five default bad channels [T7, T8, Fp1, Fp2, Tp7] and additional visual inspection: median = 1.5 channels). Data was high-pass filtered (0.01Hz), then line noise and its harmonics were removed using a DFT filter (50, 100, 150Hz). Data was segmented into trials ranging from 0.8 s before to 1.2 s after search display onset. An ICA was performed to remove oculomotor artifacts, and trials with other motor artifacts were removed from further EEG analysis as per visual inspection (*mean* = 6.49 %). Baseline correction was performed by averaging (and then subtracting from the signal) a window ranging from 0.8 s to 0.2 s before the onset of the search display.

### Eye-tracking analysis

The time window of interest spanned 1.2 seconds following the onset of the search display. Due to head movements after eye calibration, one participant had one block excluded, and another participant had two blocks excluded before eye tracking data analysis. Blink correction was performed using custom code adapted from Hershman et al. (2018). To ensure that EEG responses to the frequency-tagged stimuli (target and distractor) were not confounded by large eye movements, we implemented eye movement exclusion criteria. A circular region of interest (ROI) with a radius of 3.5 dva (no stimuli was presented at this area) was defined around the central fixation point. Trials in which participants’ gaze deviated outside this ROI for more than 50 ms were classified as saccade trials and excluded from further EEG analysis. Participants with more than 50% of their trials marked as saccade trials were excluded from group-level analyses. As a result, three participants were replaced. On average, 13.65% of trials per participant were removed based on the saccade criterion.

### RIFT responses

To quantify the degree to which the EEG signal reflects the tagging signals, we computed magnitude-squared coherence (Arora et al., 2025; Pan et al., 2021), a dimensionless measure ranging from 0 to 1 that reflects the consistency of two signals in both magnitude and phase. Coherence was computed between a reference sinusoid (sampled at 2048 Hz) and the neural responses to the tagged stimuli, separately for each frequency, EEG channel and participant. To calculate coherence for a specific frequency of interest, segmented trials (*N*) were bandpass filtered (±1.9 Hz) around the respective tagging frequency using a two-pass, fourth-order Butterworth filter with a Hamming taper. The filtered time-series data were then subjected to a Hilbert transform to extract the instantaneous magnitude (*M(t)*) and phase (*ϕ(t)*) of the signal. The set of all instantaneous magnitudes of the filtered responses 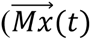 and the reference sinusoid 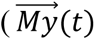 across all n trials, as well as the differences between their instantaneous phases across all n trials 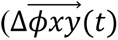 were used to compute time-varying coherence (Formula 1). Notably, when calculating coherence at 60 Hz, individual EEG trials were first phase (re)-aligned by temporally shifting them by a maximum of 16.67 ms (34 samples in EEG) based on the phase at which they were presented. This ensured accurate estimation of coherence and causes only to minimal temporal smearing (see Tagging manipulation section, above).

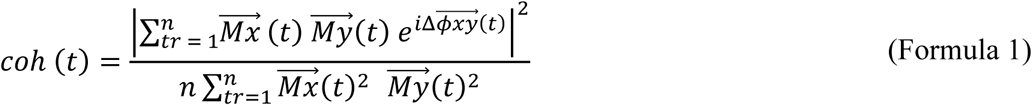

For each participant, six channels were selected based on the highest coherence averaged across the two tagging frequencies during the 1.2 s following search display onset (Arora et al., 2025; Hustá et al., 2025). Notably, previous studies have shown that the exact number of top channels selected does not substantially affect the results (Arora et al., 2025). To account for the fact that frequency-tagged stimuli evoke spatially specific neural responses that vary depending on their location on the screen, channel selection was performed separately for each of the 6 tagging locations (Minarik et al., 2023; see scalp topographies for the six locations in the supplementary materials; Figure S5). Coherence traces were then averaged across the top six selected channels, six tagging locations, and two tagging frequencies to produce a single coherence trace per condition for each participant, which was used for all subsequent EEG analyses. Coherence spectrograms (Figure 1C & 1D) were computed across frequencies from 56.8 Hz to 67.2 Hz in 0.8 Hz steps.

To examine the trial-wise correlation between RIFT responses and behavioral performance (i.e., within participants) we calculated Trial-Ensemble Phase Similarity (TEPS) as a single trial measure of RIFT. TEPS quantifies the phase similarity between each individual trial and the average phase of all other trials, using a leave-one-trial-out approach. This measure ranges from 1 (perfect alignment) to –1 (perfect opposition), capturing how closely a trial’s phase follows the group-level phase dynamics over time (see Formula 2). Specifically, for each trial *n* and time point *t*, we extracted the instantaneous phase *ϕ*_*K*_(*t*) from the bandpass-filtered EEG signal (±1.9 Hz) via the Hilbert transform. We then calculated the circular mean phase across all other trials 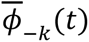 and defined TEPS as the cosine of the phase difference between *ϕ*_*K*_(*t*) and 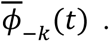

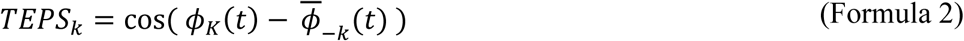

### Statistical Analysis

For the behavioral analysis, we excluded trials with reaction times shorter than 200 ms and used a paired-sample t-test to compare the mean response times (RTs) across conditions.

To statistically compare coherence traces between conditions, individual traces were subtracted between conditions and resulting individual coherence-difference traces were subjected to a cluster-based permutation test (Maris & Oostenveld, 2007). First, one-sample t-tests at the group level were performed for each time point to identify clusters where coherence traces differed significantly from zero (*p* < 0.05). Individual clusters were defined as one or more consecutive significant time points. For each cluster, the sum of t-values across all included timepoints (i.e., the t-mass) was computed and used as the cluster-level statistic. A null distribution of t-mass values was created by flipping the sign of a random selection of difference traces across 10,000 permutations and repeating the cluster identification procedure described above, except for only including the largest cluster in the null-distribution. Observed clusters were considered statistically significant if their t-mass exceeded the 95th percentile of t-masses within the null distribution.

To statistically assess correlations between neural responses in early visual cortex and behavioral performance, we computed trial-wise Pearson correlations between single trial RIFT responses (TEPS) and response times (including only trials with correct responses). We normalized the TEPS value for each location by subtracting the mean TEPS value at that location from the raw TEPS value for each trial, in order to eliminate differences in RIFT responses across locations (Figure S4). For each participant, we calculated the correlation between TEPS at each time point and RTs between trials, resulting in a time-resolved series of correlations. These time-series were assessed using the non-parametric cluster-based permutation test procedure described in the previous paragraph.

## Data and Code Availability

The raw data and analysis code for this experiment is publicly available at https://osf.io/szuxa/.

## Notes

https://osf.io/szuxa/

### Competing Interest Statement

The authors have declared no competing interest.

### Funding

- China Scholarship Council: 202308510050
- European Research Council (ERC) advanced grant: 833029 - [LEARNATTEND]
- NWO Open competition grant: 25 406.21.GO.034

## References

Arora, K., Gayet, S., Kenemans, J. L., Van der Stigchel, S., & Chota, S. (2025). Dissociating external and internal attentional selection. iScience, 28(4).

Beck, D. M., & Kastner, S. (2009). Top-down and bottom-up mechanisms in biasing competition in the human brain. Vision Research, 49(10), 1154–1165.

Born, S., Kerzel, D., & Theeuwes, J. (2011). Evidence for a dissociation between the control of oculomotor capture and disengagement. Experimental Brain Research, 208(4), 621–631.

Cosman, J. D., Lowe, K. A., Zinke, W., Woodman, G. F., & Schall, J. D. (2018). Prefrontal control of visual distraction. Current Biology, 28(9), 1330–1330.

de Fockert, J. W., & Theeuwes, J. (2012). Role of frontal cortex in attentional capture by singleton distractors. Brain and Cognition, 80(3), 367–373.

Desimone, R., & Duncan, J. (1995). Neural mechanisms of selective visual attention. Annual Review of Neuroscience, 18(1), 193–222.

Dietz, L., Strauch, C., Arora, K., Van der Stigchel, S., Chota, S., & Gayet, S. (2025). Anticipated relevance prepares visual processing for efficient memory-guided selection. bioRxiv.

Drijvers, L., Jensen, O., & Spaak, E. (2021). Rapid invisible frequency tagging reveals nonlinear integration of auditory and visual information. Human Brain Mapping, 42(4), 1138–1152.

Duecker, K., Gutteling, T. P., Herrmann, C. S., & Jensen, O. (2021). No evidence for entrainment: Endogenous gamma oscillations and rhythmic flicker responses coexist in visual cortex. Journal of Neuroscience, 41(31), 6684–6698.

Duecker, K., Shapiro, K. L., Hanslmayr, S., Griffiths, B. J., Pan, Y., Wolfe, J. M., & Jensen, O. (2025). Guided visual search is associated with target boosting and distractor suppression in early visual cortex. Communications Biology, 8(1), 912.

Duncan, D. H., Forschack, N., van Moorselaar, D., Müller, M. M., & Theeuwes, J. (2025). Learning modulates early encephalographic responses to distracting stimuli: a combined SSVEP and ERP study. Journal of Neuroscience, 45(21).

Faul, F., Erdfelder, E., Lang, A. G., & Buchner, A. (2007). G*Power 3: A flexible statistical power analysis program for the social, behavioral, and biomedical sciences. Behavior Research Methods, 39(2), 175–191.

Ferrante, O., Zhigalov, A., Hickey, C., & Jensen, O. (2023). Statistical learning of distractor suppression downregulates prestimulus neural excitability in early visual cortex. Journal of Neuroscience, 43(12), 2190–2198.

Folk, C. L., & Remington, R. (1998). Selectivity in distraction by irrelevant featural singletons: Evidence for two forms of attentional capture. Journal of Experimental Psychology: Human Perception and Performance, 24(3), 847–858.

Forschack, N., Gundlach, C., Hillyard, S., & Müller, M. M. (2022). Dynamics of attentional allocation to targets and distractors during visual search. NeuroImage, 264, 119759.

Geng, J. J. (2014). Attentional mechanisms of distractor suppression. Current Directions in Psychological Science, 23(2), 147–153.

Hershman, R., Henik, A., & Cohen, N. (2018). A novel blink detection method based on pupillometry noise. Behavior research methods, 50(1), 107–114.

Hickey, C., McDonald, J. J., & Theeuwes, J. (2006). Electrophysiological evidence of the capture of visual attention. Journal of Cognitive Neuroscience, 18(4), 604–613.

Hickey, C., Van Zoest, W., & Theeuwes, J. (2010). The time course of exogenous and endogenous control of covert attention. Experimental Brain Research, 201(6), 789–796.

Hustá, C., Meyer, A., & Drijvers, L. (2025). Using rapid invisible frequency tagging (RIFT) to probe the neural interaction between representations of speech planning and comprehension. Neurobiology of Language, 1–16.

Kleiner, M., Brainard, D., Pelli, D., Ingling, A., Murray, R., and Broussard, C. (2007). What’s new in psychtoolbox-3. Perception 36, 1–16.

Klink, P. C., Teeuwen, R. R. M., Lorteije, J. A. M., & Roelfsema, P. R. (2023). Inversion of pop-out for a distracting feature dimension in monkey visual cortex. Proceedings of the National Academy of Sciences of the United States of America, 120(9), e2210839120.

Landis, C. (1954). Determinants of the critical flicker-fusion threshold. Physiological Reviews, 34(2), 259–286.

Lin, R., Meng, X., Chen, F., Li, X., Jensen, O., Theeuwes, J., & Wang, B. (2024). Neural evidence for attentional capture by salient distractors. Nature Human Behaviour, 8(5), 932–944.

Luck, S. J., Chelazzi, L., Hillyard, S. A., & Desimone, R. (1997). Neural mechanisms of spatial selective attention in areas V1, V2, and V4 of macaque visual cortex. Journal of Neurophysiology, 77(1), 24–42.

Luck, S. J., Gaspelin, N., Folk, C. L., Remington, R. W., & Theeuwes, J. (2021). Progress toward resolving the attentional capture debate. Visual Cognition, 29(1), 1–21.

Maris, E., & Oostenveld, R. (2007). Nonparametric statistical testing of EEG and MEG data. Journal of Neuroscience Methods, 164(1), 177–190.

Minarik, T., Berger, B., & Jensen, O. (2023). Optimal parameters for rapid (invisible) frequency tagging using MEG. NeuroImage, 281, 120389.

Oostenveld, R., Fries, P., Maris, E., & Schoffelen, J. M. (2011). FieldTrip: Open source software for advanced analysis of MEG, EEG, and invasive electrophysiological data. Computational Intelligence and Neuroscience, 2011(1), 156869.

Pan, Y., Frisson, S., & Jensen, O. (2021). Neural evidence for lexical parafoveal processing. Nature Communications, 12(1), 5234.

Reynolds, J. H., & Desimone, R. (2003). Interacting roles of attention and visual salience in V4. Neuron, 37(5), 853–863.

Schneider, M., Tzanou, A., Uran, C., & Vinck, M. (2023). Cell-type-specific propagation of visual flicker. Cell Reports, 42(5), 112492.

Schubö, A. (2009). Salience detection and attentional capture. Psychological Research, 73(2), 233– 243.

Seijdel, N., Marshall, T. R., & Drijvers, L. (2023). Rapid invisible frequency tagging (RIFT): A promising technique to study neural and cognitive processing using naturalistic paradigms. Cerebral Cortex, 33(5), 1626–1629.

Spaak, E., Bouwkamp, F. G., & de Lange, F. P. (2024). Perceptual foundation and extension to phase tagging for rapid invisible frequency tagging (RIFT). Imaging Neuroscience, 2, 1–14.

Theeuwes, J. (1991). Cross-dimensional perceptual selectivity. Perception & psychophysics, 50(2), 184–193.

Theeuwes, J. (1992). Perceptual selectivity for color and form. Perception & Psychophysics, 51(6), 599–606.

Theeuwes, J. (2010). Top–down and bottom–up control of visual selection. Acta Psychologica, 135(2), 77–99.

Theeuwes, J. (2021). Response to commentaries to Luck et al. (2021): Progress toward resolving the attentional capture debate. Visual Cognition, 29(9), 637–643.

Theeuwes, J. (2025). Attentional capture and control. Annual Review of Psychology, 76, 1–24.

Theeuwes, J., Atchley, P., & Kramer, A. F. (2000). On the time course of top-down and bottom-up control of visual attention. Attention and Performance, 18, 104–124

Tsotsos, J. K., Culhane, S. M., Wai, W. Y. K., Lai, Y., Davis, N., & Nuflo, F. (1995). Modeling visual attention via selective tuning. Artificial Intelligence, 78(1–2), 507–545.

Wandell, B. A., Dumoulin, S. O., & Brewer, A. A. (2007). Visual field maps in human cortex. Neuron, 56(2), 366–383.

Wang, B., van Driel, J., Ort, E., & Theeuwes, J. (2019). Anticipatory distractor suppression elicited by statistical regularities in visual search. Journal of Cognitive Neuroscience, 31(10), 1535– 1548.

Zhigalov, A., & Jensen, O. (2020). Alpha oscillations do not implement gain control in early visual cortex but rather gating in parieto-occipital regions. Human Brain Mapping, 41(18), 5176– 5186.

Zhigalov, A., Herring, J. D., Herpers, J., Bergmann, T. O., & Jensen, O. (2019). Probing cortical excitability using rapid frequency tagging. NeuroImage, 195, 59–66.

