## Supplementary for "Dynamic competition between bottom-up saliency and top-down goals in early visual cortex"

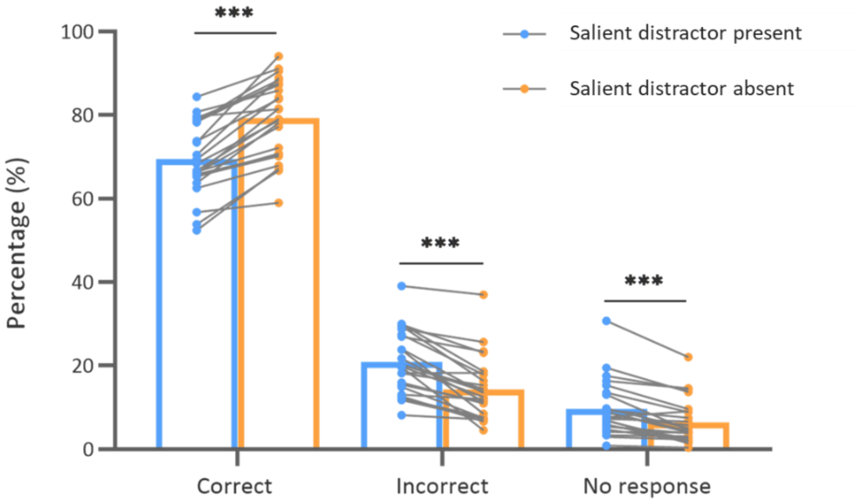


**Figure S1.** Supplemental behavioral results. Behavioral responses could be divided into correct, incorrect or no responses (within the 1.3s deadline). Participants showed a lower correct response rate, along with higher incorrect response and no response rates, when a salient distractor was present rather than absent (thus mirroring the RT results reported in the main manuscript). *** $p < 0.001$.

### A Individual coherence spectrograms (60Hz)

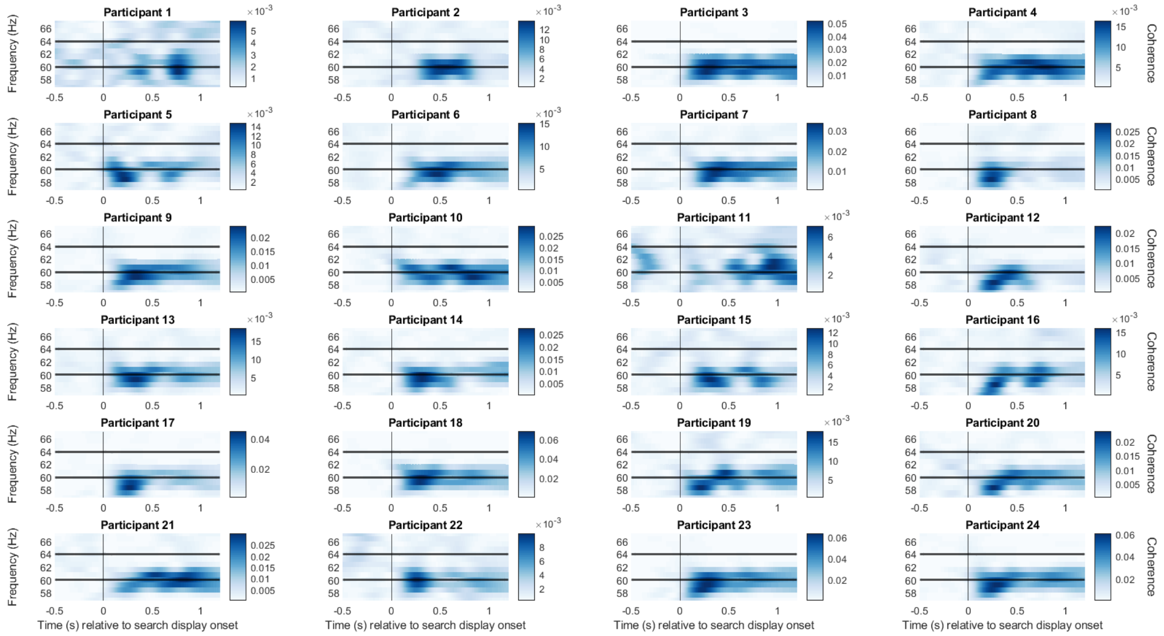


### B Individual scalp topography (60 Hz)

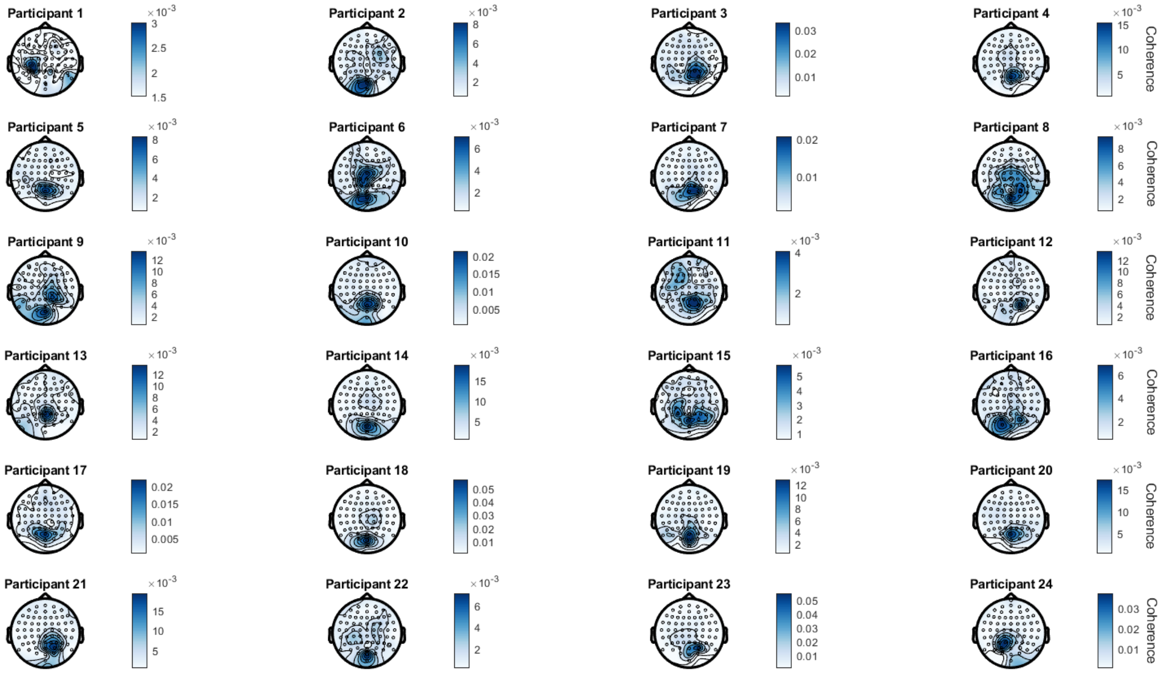


**Figure S2. (A)** Coherence spectrograms at 60 Hz for each participant, showing a clear peak at 60 Hz following flicker onset. **(B)** Scalp topographies for each participant, depicting average 60 Hz coherence across the 1.2-second search display. Strongest responses were observed over parieto-occipital electrodes.

**A** Individual coherence spectrograms (64Hz)

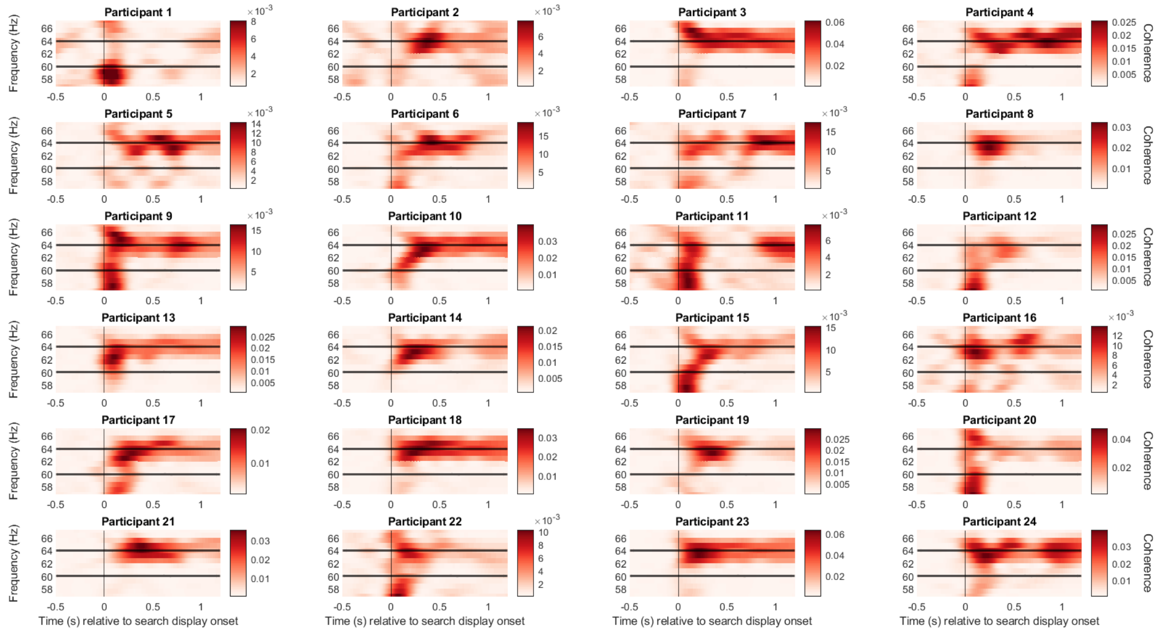


**B** Individual scalp topography (64Hz)

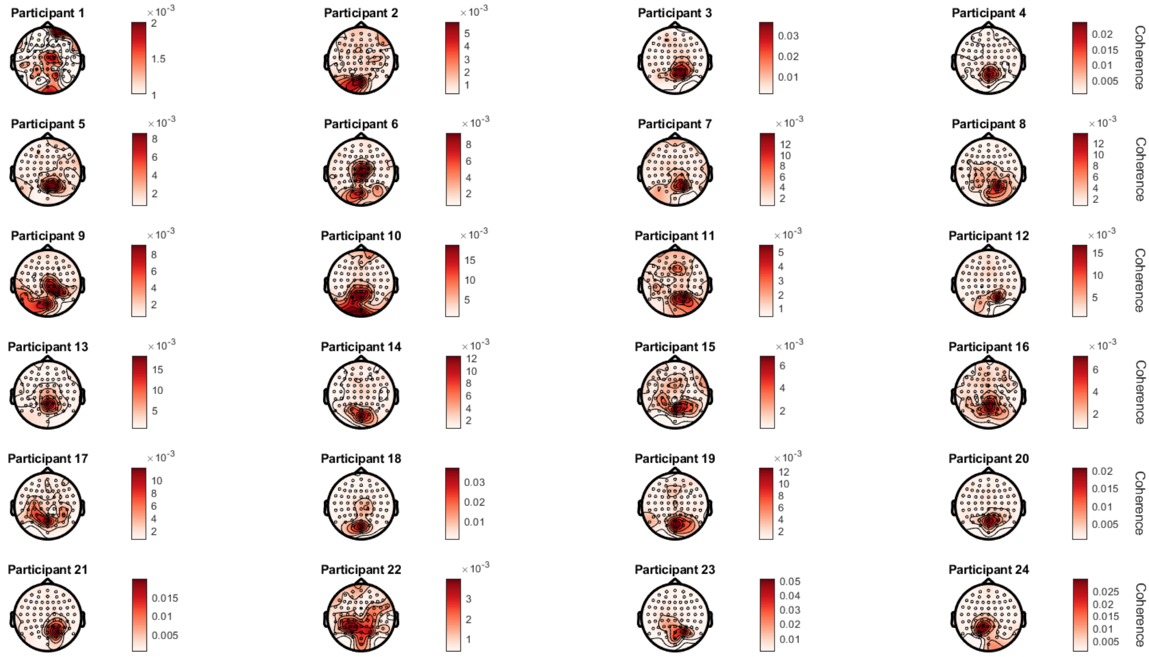


**Figure S3. (A)** Coherence spectrograms at 64 Hz for each participant, showing a clear peak at 64 Hz following flicker onset. **(B)** Scalp topographies for each participant, depicting average 64 Hz coherence across the 1.2-second search display. Strongest responses were observed over parieto-occipital electrodes.

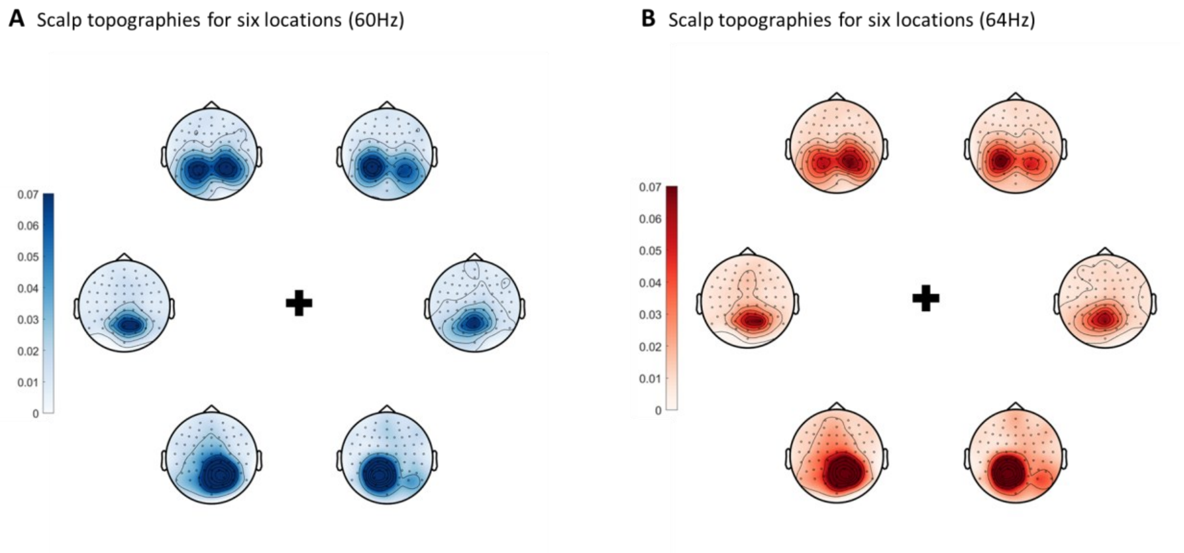


**Figure S4.** Scalp topographies corresponding to six stimulus locations on search display for **(A)** 60 Hz and **(B)** 64 Hz frequency tagging. The maps show average coherence over the 1.2-second search display, indicating that frequency-tagged stimuli evoked spatially specific neural responses.

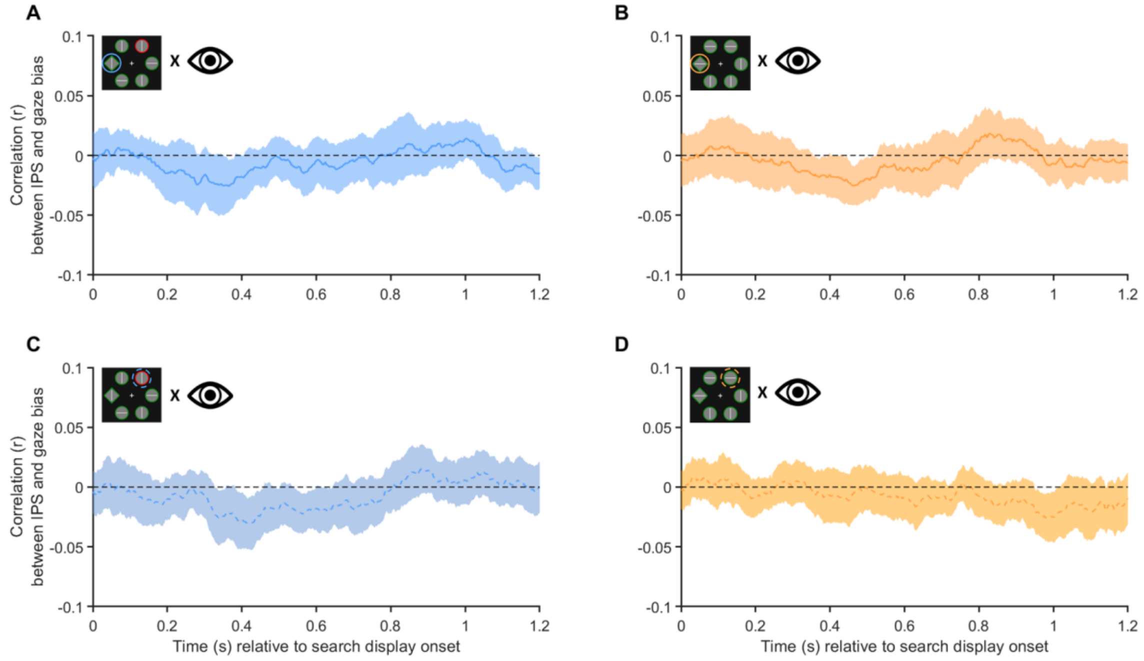


**Figure S5.** Time-resolved, trial-wise correlation between gaze bias and IPS responses of the **(A)** target with salient distractor, of the **(B)** target without salient distractor, of the **(C)** salient distractor, and of the **(D)** tagged non-salient distractor. Gaze bias refers to the distance between the gaze position and the center of the tagged stimulus (i.e., target or distractor). Shaded areas represent the 95% confidence intervals of the mean. Thus, gaze bias did not reliably predict RIFT responses.
